# Ethanol consumption mediates parasitoid resistance via effects on host metabolism

**DOI:** 10.1101/2025.10.01.679850

**Authors:** Carrie L. Marean-Reardon, Patrick N. Reardon, Nathan T. Mortimer

## Abstract

Many insect species use self-medication, the consumption of an environmental compound with antipathogen activity, as a defense against pathogen infection. One well-studied example is the interaction between *Drosophila melanogaster* and parasitoid wasps, in which *D. melanogaster* larvae consume ethanol-laden food to kill the developing parasitoid. Despite research into self-medication as a behavioral immune response to parasitoid infection, the parasitoid-killing mechanism remains elusive. To test the impact of ethanol consumption and infection on host metabolism, we used untargeted Nuclear Magnetic Resonance (NMR) metabolomics of hemolymph samples isolated from naïve and parasitoid infected larvae fed ethanol-containing or control food. Surprisingly, we found that the consumption of dietary ethanol did not result in an elevated hemolymph ethanol abundance. Instead, we found evidence that host carbohydrate-derived energy production and amino acid metabolism were altered by ethanol consumption. Our results suggest that these ethanol-mediated changes in host metabolism, rather than a direct effect of dietary ethanol, confers parasitoid resistance.

## Introduction

Drosophilid species are commonly infected by parasitoid wasps, and following infection, fly hosts may mount either an immune-mediated or a behavioral defense response (Mortimer & Schlenke, 2025). The major immune response against parasitoid infection is the hemocyte-mediated encapsulation and melanization of the parasitoid egg (Mortimer & Schlenke, 2025). This immune response is supplemented by several behavioral defenses, including self-medication, in which infected individuals alter their feeding behavior to consume specific compounds with antiparasitoid properties, which they would otherwise avoid or consume in smaller quantities (Abbott, 2014; Mortimer & Schlenke, 2025). A medication defense response to infection is seen in parasitoid-infected *Drosophila melanogaster* (Milan et al., 2012) and *Drosophila suzukii* (Poyet et al., 2017), and more broadly across insects including Lepidopteran and Hymenopteran species (Abbott, 2014).

The consumption of environmental ethanol by parasitoid-infected *D. melanogaster* larvae is a well-studied form of self-medication (Milan et al., 2012; Lynch et al., 2017). In this interaction, the parasitoid lays its egg directly into the larval hemocoel, triggering a cellular immune response in which the parasitoid is encapsulated and melanized (Mortimer & Schlenke, 2025). If the parasitoid is able to evade host immunity, the wasp egg hatches into a larva which develops in the host hemolymph environment (Carton et al., 1986). Host medication via ethanol consumption increases the likelihood of fly survival following infection, however it does so independently of the cellular immune response (Lynch et al., 2016). This suggests that medication causes a change in the hemolymph composition so that it is either toxic to the parasitoid or lacking an essential nutrient for parasitoid survival.

It has been proposed that fly ethanol consumption results in a subsequent increase in systemic ethanol within their hemolymph, and that this ethanol acts as a toxin leading to parasitoid death (Sadanandappa et al., 2024). This model assumes that dietary ethanol passes directly from the gut into the host hemolymph following consumption. We propose an alternate model in which ethanol consumption alters *D. melanogaster* metabolism, bringing about changes in the hemolymph metabolite pool that have an antiparasitoid effect. These putative models underlying host medication will be assessed through the characterization of *D. melanogaster* hemolymph metabolites in naïve and infected larvae fed distinct diets.

## Materials and Methods

### Fly maintenance and husbandry

Oregon R stock (RRID: BDSC_25211) were maintained at room temperature. Adult flies (25 female, 15 male) were selected and allowed to lay on yeast smeared grape juice plates in embryo cages. Cages were maintained in a 24°C incubator with a 12-hour light/dark cycle. The plate was changed every 24 hours and was allowed to age an additional 24 hours to maximize the number of 2^nd^ instar larvae available.

### Parasitoid and host survival assay

2^nd^ instar larvae were transferred to vials of freshly prepared Formula 4-24 Instant Drosophila Medium, Blue food (Carolina Biological Supply #173210) with no additives (control) or 6% ethanol. Uninfected larvae were allowed to mature; larvae in the infection groups were allowed 24 hours to acclimate to the food before being infected. For infection vials, *Leptopilina boulardi* or *Leptopilina heterotoma* wasps (6-8 females, 3-4 males) were introduced, given 6 hours to infect, and then removed and fly larvae were allowed to mature without further interference. Pupae were tallied upon emergence, and flies and wasps were tallied upon eclosion. The experiments were maintained in a 24°C incubator with 12-hour light/dark cycle.

### Hemolymph preparation for NMR

2^nd^ instar larvae were transferred to plates of freshly prepared Formula 4-24 Instant Drosophila Medium, Blue food (Carolina Biological Supply #173210) with no additives (control) or 6% ethanol. Naïve plates were allowed to grow an additional 72 hours at 24°C before hemolymph collection. For infection plates, larvae were allowed to acclimate to the food overnight before infection. *Leptopilina boulardi* or *Leptopilina heterotoma* wasps (6-8 females, 3-4 males) were introduced, given 6 hours to infect, and then removed and larvae were allowed 42 hours to further mature. Larvae were collected as close as possible to fat wandering 3^rd^ instars to maximize hemolymph volume, and lightly mashed about the mouth-hooks to release hemolymph. Harvested larva were spun at 5000xg for 2 minutes in hemolymph collection devices. Collection tubes were weighed before and after collection of hemolymph to normalize calculations with mass of hemolymph. Hemolymph was then mixed with 160 ul ddH2O, 20 ul 10x PBS pH 7.6, and 20 ul IS-2 Chenomx Internal Standard. Twelve larvae were used for each sample, with 3-5 replicates per sample. Samples were either frozen at -80C for later NMR spectra collection or collected that day. Hemolymph collection devices were made by cutting a thin slit, with a razor blade, across the bottom of a 600ul Eppendorf tube and placing that tube in a 1.6 ml Eppendorf tube. Blue food recipe ratio was 0.2g food per 1 ml dH2O. 35mm plates received 3.6g hydrated food, 60 mm plates received 9g food, and each vial received 4.5 g food. Ethanol plates were prepared similarly except 6% v/v ethanol was added after the food was hydrated, as the ethanol affected hydration if added with the water.

### NMR metabolomics and profiling

Samples were placed in Bruker 3mm NMR tubes (Bruker # Z172600). NMR data were collected on an 800 MHz Bruker Avance 3 HD equipped with a 5mm TCI (HCN) cryogenic probe. 1D 1H data were collected using the Chenomx recommended pulse sequence (noesyphpr). NMR experimental parameters were set to the following Chenomx recommended values, spectral window of 12 ppm, acquisition time of 4s, recycle delay of 1s and 512 scans. All NMR experiments were performed at 25 C. Data were apodized using 0.5 Hz line broadening, zero filled to twice the size, Fourier Transformed, phased, and baseline corrected using the Chenomx NMR Suite. NMR data were profiled using the Chenomx NMR Suite to identify and quantify the metabolites.

### Statistical analysis

All statistical analysis was performed in R (R Core Team, 2025), and plots were generated using the *ggplot2* R package (Wickham, 2016). Eclosion success data were compared by Analysis of Variance of Aligned Rank Transformed Data followed by Dunnett’s test using the *ARTool* and *multcomp* packages in R (Hothorn et al., 2008; Wobbrock et al., 2011). Metabolomics data were normalized by Probability Quotient Normalization (PQN) using the *Rcpm* package in R (Derks, 2016), and tested for normality using the *nortest* package in R (Gross & Ligges, 2015). Any metabolites with a non-normal distribution were log_10_ transformed. Data were converted to Z scores to allow for direct comparison. Metabolite abundances were compared by Welch Two Sample t-test for the contrasts indicated in the text.

## Data availability

The data underlying this study are available from Open Science Framework at https://osf.io/ptqzf/?view_only=75e893c66c074d8ca18e7c3f4545c1b3.

## Results

In line with previous findings, we found that consumption of food containing 6% ethanol acted as a medication defense against parasitoid infection in *D. melanogaster* larvae. We found that while feeding on 6% ethanol food had a deleterious effect on naïve *D. melanogaster* larvae (Figure 1A; p = 2.3×10^-4^), ethanol consumption significantly decreased the success of the parasitoid *Leptopilina heterotoma*, with the proportion of larvae eclosing as adult wasps reduced from 76.4% on control food to 19.6% on 6% ethanol food (Figure 1B; p = 0.001). Interestingly, this medication effect has been demonstrated to be less effective against specialist parasitoid species (Lynch et al., 2017), and our data support this conclusion, with ethanol consumption exerting a marginal effect on the survival of the specialist parasitoid *Leptopilina boulardi* (Figure 1B; p = 0.064).

**Figure 1.**
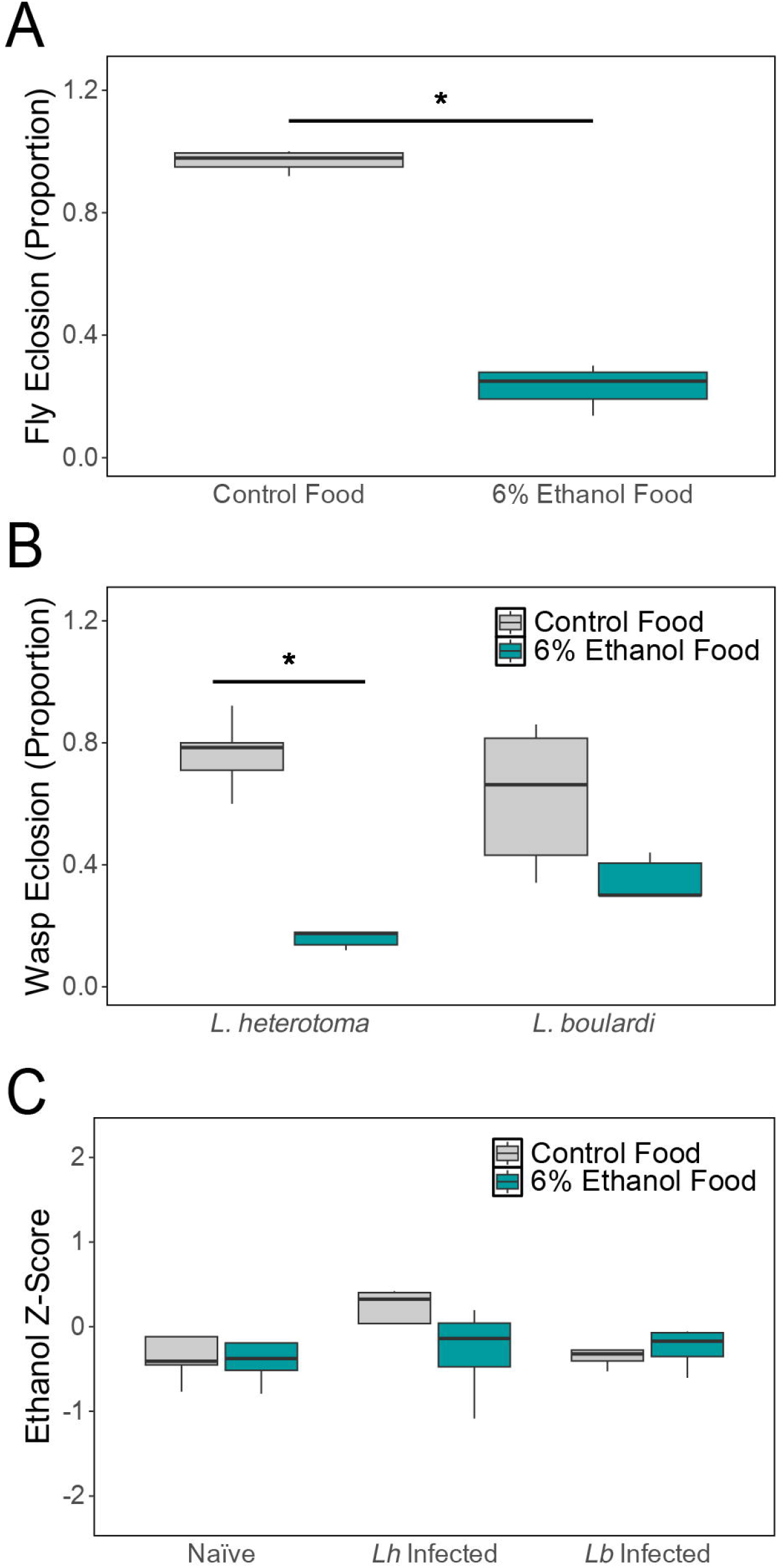
Box-whisker plots of the proportion of larvae successfully eclosing as adult flies from naïve larvae (A), or adult wasps from infected larvae (B). * indicates p <0.05 by Dunnett’s test. Larvae fed control food are shown in gray, larvae fed 6% ethanol food are shown in dark cyan. (C) Box-whisker plot of hemolymph ethanol abundance by Z score in the indicated samples. Samples derived from larvae fed control food are shown in gray, and from larvae fed 6% ethanol food are shown in dark cyan.

To test the hypothesis that systemic ethanol is the molecule leading to parasitoid death, we isolated hemolymph from larvae consuming control and 6% ethanol-laden food and used NMR to quantify hemolymph ethanol abundance. While we detected ethanol in all of our hemolymph samples, the hemolymph from larvae fed on 6% ethanol containing food showed no increase in ethanol abundance (Figure 1C; ΔZ = -0.18, p = 0.75). Similarly, there was no increase in hemolymph ethanol content in parasitoid-infected larvae fed on 6% ethanol containing food when compared to infected larvae on control food (Figure 1C; *L. heterotoma* infection: ΔZ = -0.41, p = 0.31; *L. boulardi* infection: ΔZ = 0.11, p = 0.48), suggesting that hemolymph ethanol content is not linked to infection or host immunity.

Despite this surprising result, we detected significant changes in the abundance of other hemolymph metabolites following ethanol consumption in naïve larvae. Identifying these changes can provide insight into changes in host physiology and metabolism following ethanol consumption and define the putative antiparasitoid hemolymph condition. We found that naïve larvae fed 6% ethanol food had a decreased hemolymph abundance of lactate (Figure 2A; ΔZ = -1.42, p = 0.04), citrate (Figure 2B; ΔZ = -3.19, p = 0.01), and 2-hydroxyglutarate (Figure 2C; ΔZ = -1.43, p = 0.03). These metabolites are linked to the production of cellular energy via glycolysis and the tricarboxylic acid (TCA) cycle, suggesting that this process is altered by ethanol consumption. The decrease in hemolymph levels of these metabolites likely corresponds to an increased demand for energy production in host cells. This is further supported by a marginal decrease in circulating trehalose (ΔZ = -2.64, p = 0.09), the primary source of sugar for energy production in *D. melanogaster* (Reyes-DelaTorre et al., 2012). Alongside these possible changes in cellular energy production, we found altered hemolymph abundance of the amino acids aspartate (Figure 2D; ΔZ = 4.68, p = 0.003), arginine (Figure 2E; ΔZ = 2.14, p = 0.006), and threonine (Figure 2F; ΔZ = -2.54, p = 0.008), suggesting that amino acid metabolism or protein stability are sensitive to ethanol consumption. Interestingly, aspartate synthesis is linked to the TCA cycle. Our observation that ethanol consumption increases levels of aspartate, while decreasing citrate abundance may suggest that citrate synthase activity is altered in ethanol fed larvae. Finally, we found a decreased abundance of trimethylamine (TMA) in the hemolymph of ethanol consuming larvae (Figure 2G; ΔZ = -2.57, p = 0.002). TMA Is a byproduct of gut microbiome metabolism (Falony et al., 2015), suggesting that dietary ethanol may also alter the microbiome community function.

**Figure 2.**
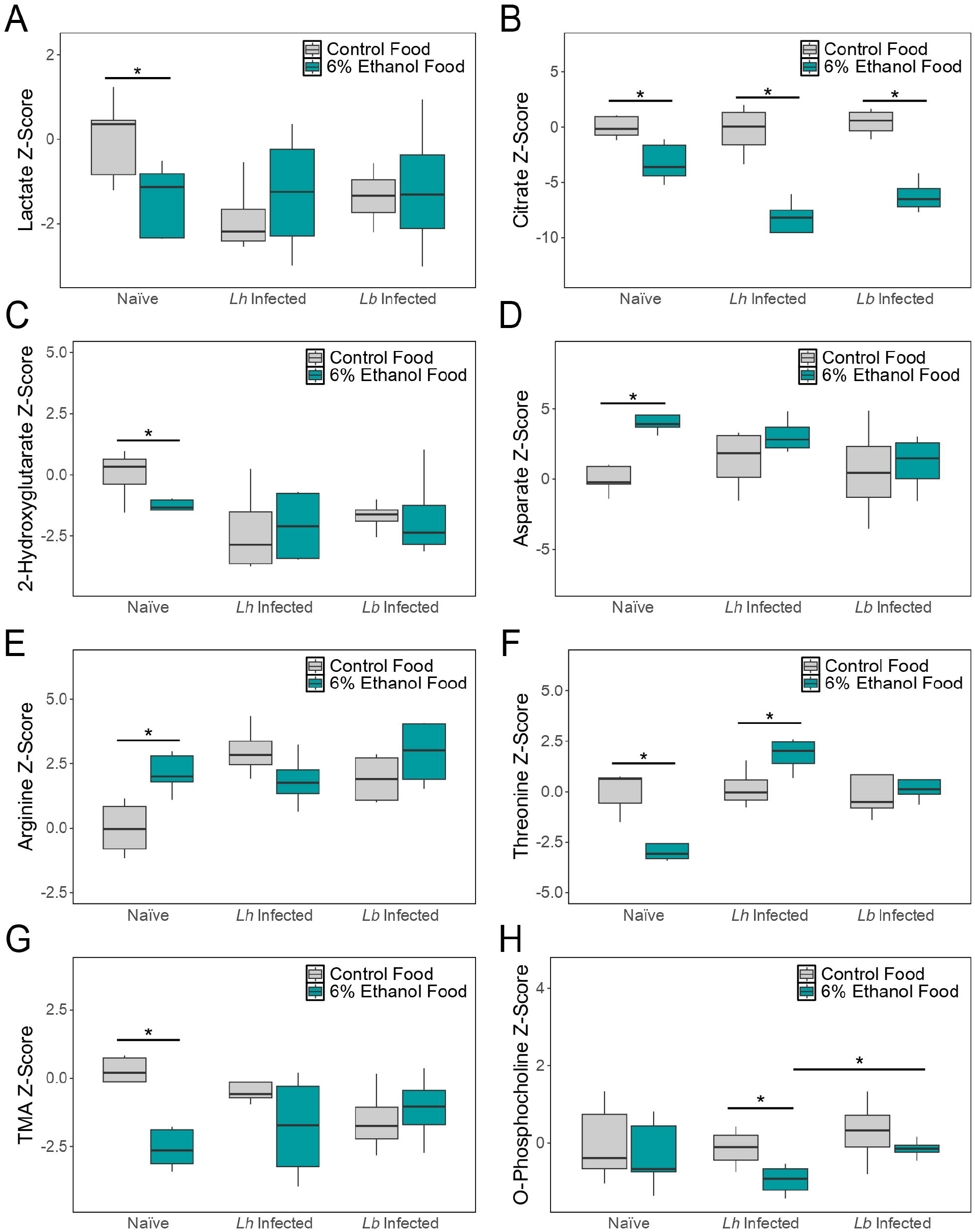
Box-whisker plots of hemolymph abundance by Z score in the indicated samples; (A) lactate, (B) citrate, (C) 2-hydroxyglutarate, (D) aspartate, (E) arginine, (F) threonine, (G) TMA, and (H) O-phosphocholine. * indicates p <0.05 by Welch Two Sample t-test. In all panels, samples derived from larvae fed control food are shown in gray, and from larvae fed 6% ethanol food are shown in dark cyan.

We found that many of these ethanol consumption-induced changes in larval hemolymph metabolites were specific to naïve larvae. Only citrate (Figure 2B) was consistently altered by ethanol consumption in naïve, *L. heterotoma-* and *L. boulardi*-infected fly larvae (*L. heterotoma* infection: ΔZ = -8.56, p = 0.004; *L. boulardi* infection: ΔZ = -6.65, p = 6.4×10^-4^). The amino acid threonine, which was decreased in ethanol fed naïve larvae, was instead increased in abundance in the hemolymph of ethanol fed *L. heterotoma*-infected larvae (Figure 2F; *L. heterotoma* infection: ΔZ = 1.64, p = 0.048). O-phosphocholine was the only other identified metabolite to show altered abundance due to ethanol consumption in *L. heterotoma*-infected larvae (Figure 2H; ΔZ = -0.82, p = 0.043). Notably, O-phosphocholine abundance was also different between hemolymph samples from *L. heterotoma-* and *L. boulardi*-infected fly larvae following ethanol consumption (Figure 2H; ΔZ = -0.81, p = 0.019).

## Discussion

Our findings support a model in which self-medication by ethanol consumption confers protection against parasitoid infection by alterations in *D. melanogaster* metabolism, rather than via a direct effect of dietary ethanol. This antiparasitoid effect could either be due to an excess of a toxic metabolite within the hemolymph, or due to the depletion of an essential nutrient or inhibition of a host metabolic process that is required for parasitoid development. We identified several ethanol-responsive metabolites that may act to disrupt parasitoid development within the hemolymph. Future testing will be required to determine their roles in medication-mediated parasitoid resistance.

We also observed differences in metabolite abundance between naïve and infected larvae, regardless of food source. This suggests that the changes in certain metabolites may come from infection-induced host mechanisms or from the activity of parasitoid venom proteins. Parasitoid wasps have evolved venom proteins that suppress host immunity, and the venoms of both *L. heterotoma* and *L. boulardi* also contain proteins with homology to metabolic enzymes (Goecks et al., 2013). These parasitoid mediated changes may come from either inhibiting host metabolism or from hijacking host metabolites to alter their availability in the hemolymph.

Despite the minimal differences between hemolymph metabolites in *L. heterotoma*- and *L. boulardi*-infected larvae, *L. heterotoma* larvae appear more susceptible to the ethanol consumption-mediated medication response (Figure 1B) (Milan et al., 2012; Lynch et al., 2017). This could be explained through several possible models. *L. heterotoma* may be more sensitive to a common change between the infections. For instance, *L. heterotoma* may have an elevated demand for citrate during development, or may be more responsive to the energy production state of the host. Alternatively, the difference in susceptibility may be due to a unique change in hemolymph metabolites, based on parasitoid biology or venom activity. Interestingly, we see a significant change in O-phosphocholine levels in *L. heterotoma*-but not *L. boulardi*-infected larvae. Previous findings suggest that *L. heterotoma* may have decreased lipid synthesis (Visser et al., 2010), and so the lack of O-phosphocholine, an important component for phospholipid synthesis, may lead to parasitoid lethality.

Finally, it is important to consider the limitations of our study. One of the major limitations of any metabolomic study is that is not possible to detect every metabolite present in a sample with any given approach (Johnson & Gonzalez, 2012). This is due to the chemical properties of the metabolites themselves, as well as sensitivity thresholds for detection. Furthermore, additional compounds may be detected, but in the absence of reliable standards they may not be mapped for identification. Fortunately, ethanol is easily detected by NMR metabolomics, and we determined that dietary ethanol is not increasing the hemolymph ethanol concentration and therefore is not acting as a direct toxin as previously proposed. The hemolymph metabolites we detected as altered by ethanol consumption largely fell into the categories of energy production via glycolysis/TCA and amino acid metabolism. This suggests that these mechanisms are likely altered by ethanol feeding, and so are likely responsible for the pathogen-killing effect. Future studies will make use of targeted metabolomics to identify additional metabolites, before functional testing of their effect on parasitoid development.

## Author Contributions

Conceptualization: CMR,PNR,NTM; Formal analysis: CMR,PNR,NTM; Funding acquisition: NTM; Investigation: CMR; Methodology: CLM,PNR; Writing – original draft: CMR,PNR,NTM; Writing – review & editing: CLM,PNR

## Acknowledgements

This work was supported by NIH grant R35 GM133760 (NTM), and funding from Oregon State University. *Drosophila melanogaster* lines obtained from the Bloomington Drosophila Stock Center (NIH P40OD018537) were used in this study. The NMR Facility at Oregon State University was supported by the National Science Foundation MRI Grant 2320189, National Institutes of Health, HEI Grant 1S10OD018518, and by the M. J. Murdock Charitable Trust grant #2014162.

## References

Abbott J. 2014: Self-medication in insects: current evidence and future perspectives. — Ecological Entomology 39: 273–280.

Carton Y., Bouletreau M., Van Alphen J.J.M. & Van Lenteren J.C. 1986: The Drosophila parasitic wasps. The Genetics and Biology of Drosophila. Academic Press, London.

Derks R. 2016: Rcpm: Rcpm: General functions for the Center of Proteomics and Metabolomics.

Falony G., Vieira-Silva S. & Raes J. 2015: Microbiology meets big data: the case of gut microbiota-derived trimethylamine. — Annu Rev Microbiol 69: 305–321.

Goecks J., Mortimer N.T., Mobley J.A., Bowersock G.J., Taylor J. & Schlenke T.A. 2013: Integrative approach reveals composition of endoparasitoid wasp venoms. — PLOS One8: e64125.

Gross J. & Ligges U. 2015: nortest: Tests for Normality.

Hothorn T., Bretz F. & Westfall P. 2008: Simultaneous inference in general parametric models. — Biom J 50: 346–363.

Johnson C.H. & Gonzalez F.J. 2012: Challenges and opportunities of metabolomics. — J Cell Physiol 227: 2975–2981.

Lynch Z.R., Schlenke T.A., Morran L.T. & De Roode J.C. 2017: Ethanol confers differential protection against generalist and specialist parasitoids of Drosophila melanogaster. — PLoS ONE 12: e0180182.

Lynch Z.R., Schlenke T.A. & De Roode J.C. 2016: Evolution of behavioural and cellular defences against parasitoid wasps in the Drosophila melanogaster subgroup. — J. Evol. Biol. 29: 1016–1029.

Milan N.F., Kacsoh B.Z. & Schlenke T.A. 2012: Alcohol consumption as self-medication against blood-borne parasites in the fruit fly. — Curr. Biol. 22: 488–493.

Mortimer N.T. & Schlenke T.A. 2025: Multifaceted defenses against parasitoid wasps in Diptera. — Annual Review of Genetics 59.

Poyet M., Eslin P., Chabrerie O., Prud’Homme S.M., Desouhant E. & Gibert P. 2017: The invasive pest Drosophila suzukii uses trans-generational medication to resist parasitoid attack. — Sci Rep 7: 43696.

R CORE TEAM. 2025: R: A language and environment for statistical computing.

Reyes-Delatorre A., PeÑa-Rangel M.T., Riesgo-Escovar J.R., Reyes-Delatorre A., PeÑa-Rangel M.T. & Riesgo-Escovar J.R. 2012: Carbohydrate metabolism in Drosophila: Reliance on the disaccharide trehalose. Carbohydrates - Comprehensive Studies on Glycobiology and Glycotechnology. IntechOpen.

Sadanandappa M.K., Ahmad S., Mohanraj R., Ratnaparkhi M. & Sathyanarayana S.H. 2024: Defensive tactics: lessons from Drosophila. — Biol Open 13: bio061609.

Visser B., Lann C.L., Blanken F.J. DEN, Harvey J.A., Alphen J.J.M. VAN & Ellers J. 2010: Loss of lipid synthesis as an evolutionary consequence of a parasitic lifestyle. — PNAS 107: 8677–8682.

Wickham H. 2016: ggplot2: Elegant graphics for data analysis. Springer-Verlag, New York.

Wobbrock J.O., Findlater L., Gergle D. & Higgins J.J. 2011: The aligned rank transform for nonparametric factorial analyses using only anova procedures143–146.

